# Selecting Student-Authored Questions for Summative Assessments

**DOI:** 10.1101/2020.07.28.225953

**Authors:** Alice Huang, Dale Hancock, Matthew Clemson, Giselle Yeo, Dylan J Harney, Paul Denny, Gareth Denyer

## Abstract

Production of high-quality multiple-choice questions (MCQs) for both formative and summative assessment is a time-consuming task requiring great skill, creativity, and insight. The transition to online examinations, with the concomitant exposure of previously tried-and-tested MCQs, exacerbates the challenges of question production, and highlights the need for innovative solutions.

Several groups have shown that it is practical to leverage the student cohort to produce a very large number of syllabus-aligned MCQs for study banks. Although student-generated questions are well suited for formative feedback and practice activities, they are generally not thought to be suitable for high-stakes assessments. In this study, we aimed to demonstrate that training can be provided to students in a scalable fashion to generate questions of similar quality to those produced by experts, and that identification of suitable questions can be achieved with minimal academic review and editing.

Biochemistry and Molecular Biology students were assigned a series of activities designed to coach them in the art of writing and critiquing MCQs. This training resulted in the production of over one thousand MCQs that were then gauged for potential by either expert academic judgement, or via a data-driven approach in which the questions were trialled objectively in a low-stakes test. Questions selected by either method were then deployed in a high-stakes in-semester assessment alongside questions from two academically authored sources: textbook-derived MCQs, and past paper questions.

A total of 120 MCQs from these four sources were deployed in assessments attempted by over 600 students. Each question was subjected to rigorous performance analysis, including the calculation of standard metrics from classical test theory and more sophisticated Item Response Theory (IRT) measures. The results showed that MCQs authored by students and selected at low cost performed as well as questions authored by academics, illustrating the potential of this strategy for the efficient creation of large numbers of high quality MCQs for summative assessment.

## Introduction

The production of high-quality multiple-choice questions (MCQs) has always been a significant challenge for teachers. There is a constant demand for MCQs for both summative assessments and as resources for students to revise and engage with course content. The construction of appropriately aligned and pitched MCQs is not a trivial task. MCQs are often criticised for not assessing conceptual understanding and, instead, being overly focused on recall of subject-level minutiae (Biggs & Tang, 2011). Indeed, a large scale review of MCQs used across the United States in university biology courses, including molecular biology, revealed that more than 90% targeted the lowest two levels of the revised Bloom’s taxonomy (Momsen et al., 2010). However, like all forms of assessment, it is possible to write MCQs that test deep understanding, extrapolation, and other educational outcomes high on Bloom’s Taxonomy (Harper, 2003). To do this, the teacher must not only possess mastery of their subject material, but also be aware of their students’ insights and misconceptions.

The recent increased deployment of online examinations exacerbates the problem of MCQ supply and demand. Any material used in online assessments must be assumed to be in the public domain and, therefore, cannot be reused in subsequent assessments without risks to academic integrity. The danger of reusing questions on high-stakes exams has long been acknowledged (McCoubrie, 2004), with recent empirical evidence revealing considerable deterioration in the psychometric properties of questions when reused over several years (Panczyk et al., 2018). This has particular impact on ‘keeper’ questions; those in which every experienced academic curates and re-uses in exams over several years, each time reflecting on the performance metrics of the item and perhaps subtly modifying to give ever improved discrimination and power. The problem is further compounded by the fact that even the most thoughtfully constructed questions need to be validated in real assessment situations to confirm student interpretation, identify ambiguities, and validate assumptions about difficulty and discrimination. In response to the issues above, it is not surprising that many academics are attracted to use pre-prepared and presumably field-tested question banks provided by textbook publishers and in online repositories. Yet this is not a panacea, as even commercial question banks frequently contain items with flaws (Masters et al., 2001) and the questions themselves may not align well to bespoke syllabuses.

Some academics have explored the production of large pools of MCQs by their students through the use of crowdsourcing (Aflalo, 2018; Amini et al., 2020; McLeod & Snell, 1996). The resulting banks of student-generated questions are typically used for formative feedback and practice, and often prove popular resources for study and exam revision (Duret et al., 2018; Gooi & Sommerfeld, 2015; Papinczak et al., 2012; Walsh et al., 2018). A widely used tool for supporting such activities is PeerWise, an online platform where students can author and answer questions, as well as provide feedback on questions created by their peers (Denny et al., 2008; Denny, Luxton-Reilly & Hamer, 2008).

Despite growing evidence supporting the pedagogical value of getting students to create questions for each other, most instructors would be reluctant to use the student-generated questions in formal, summative assessments. The quality of the questions that students produce can vary widely (Bottomley & Denny, 2011; Purchase et al., 2010; Snow et al., 2019) and, in addition to minor deficiencies in the clarity of wording and quality of plausible distractors, both Bates et al. in Physics (Bates et al., 2014) and by Galloway and Burns in Chemistry (Galloway & Burns, 2015) found that about 5% of student-authored questions were fundamentally incorrect.

Our own observation over several years of using PeerWise to promote engagement and reflection of course learning outcomes is that students are well-placed to recognise dissonance between their own and their peers’ insights, which can reveal misconceptions by either party. We were also aware of studies in which instructors proactively improved the quality of the questions produced by students by providing the class with MCQ-writing manuals covering structural and content elements (Jobs et al., 2013), and having students attend dedicated MCQ-writing tutorials before authoring their own questions (Bates et al., 2014). We therefore hypothesized that, with adequate coaching in the art of creating MCQs, and with allocation of learning outcomes to student-authors at a suitable level of granularity, we could leverage the class to produce a large bank of assessment-grade MCQs. We further hypothesized that it would be possible to screen for the most suitable student-designed questions using performance metrics from a broadly implemented low-stakes assessment.

Accordingly, we scaffolded activities over the semester to develop skills in authoring and critiquing MCQs, specifically training students to incorporate peer-confessed insights and misconceptions into question stems and distractors. Student-authored questions were then evaluated in two different ways to assess suitability for inclusion in a high-stakes assessment: a) performing an objective, data-driven approach, by setting a low-stakes assessment and using performance data to identify questions with potential; and b) taking a hypothesis-driven approach, by identifying candidate questions through academic review and editing. Student-authored questions from both sources were then pooled with textbook-derived MCQs, and past paper questions, which have been used in previous years’ exams. The performance of all these questions were then evaluated using traditional evidence-based metrics (difficulty and discrimination index), as well as more detailed techniques (Item Response Theory and distractor analysis).

## Methods

### Ethics

Processes were conducted in accordance with the Sydney University Human Ethics protocol “Investigating how engagement with peer-generated assessment impacts student success. Project number: 2017/131”. Under this protocol, students were able to anonymously, and without prejudice, object to their contributions being included through an online form set up by the University Research Office. An independent administrator, not involved with the study, performed the cross-checking.

### Overview of Activities

Activities were organised into three cycles (Figure 1), each with the outcome of generating, or selecting for questions for a summative, high-stakes, in-semester exam – the Week 13 examination (W13E). In Cycle 1, students were trained to dissect the intent and structure of existing MCQs, gain the skills to recognise attributes of strong and weak MCQs, be empowered with the lexicon and phrasing appropriate for articulating their critiques. In Cycle 2, students were coached to author MCQs. By the end of this, each student should have written an MCQ, received peer feedback, and edited their questions according to that feedback. In Cycle 3, we determined performance metrics using a low-stakes test, and ultimately used to inform the choice of questions in the W13E. This included student-authored questions from Cycle 2 (SAMCQs), and MCQs derived from three other sources: instructor-authored past paper questions (IAPPQs), textbook-derived questions from Cycle 1 (TDQs), and student-authored, instructor-edited questions (SAIEQs), which had been created in PeerWise exercises the previous year, selected for potential, and revised by academics before deployment.

**Figure 1.**
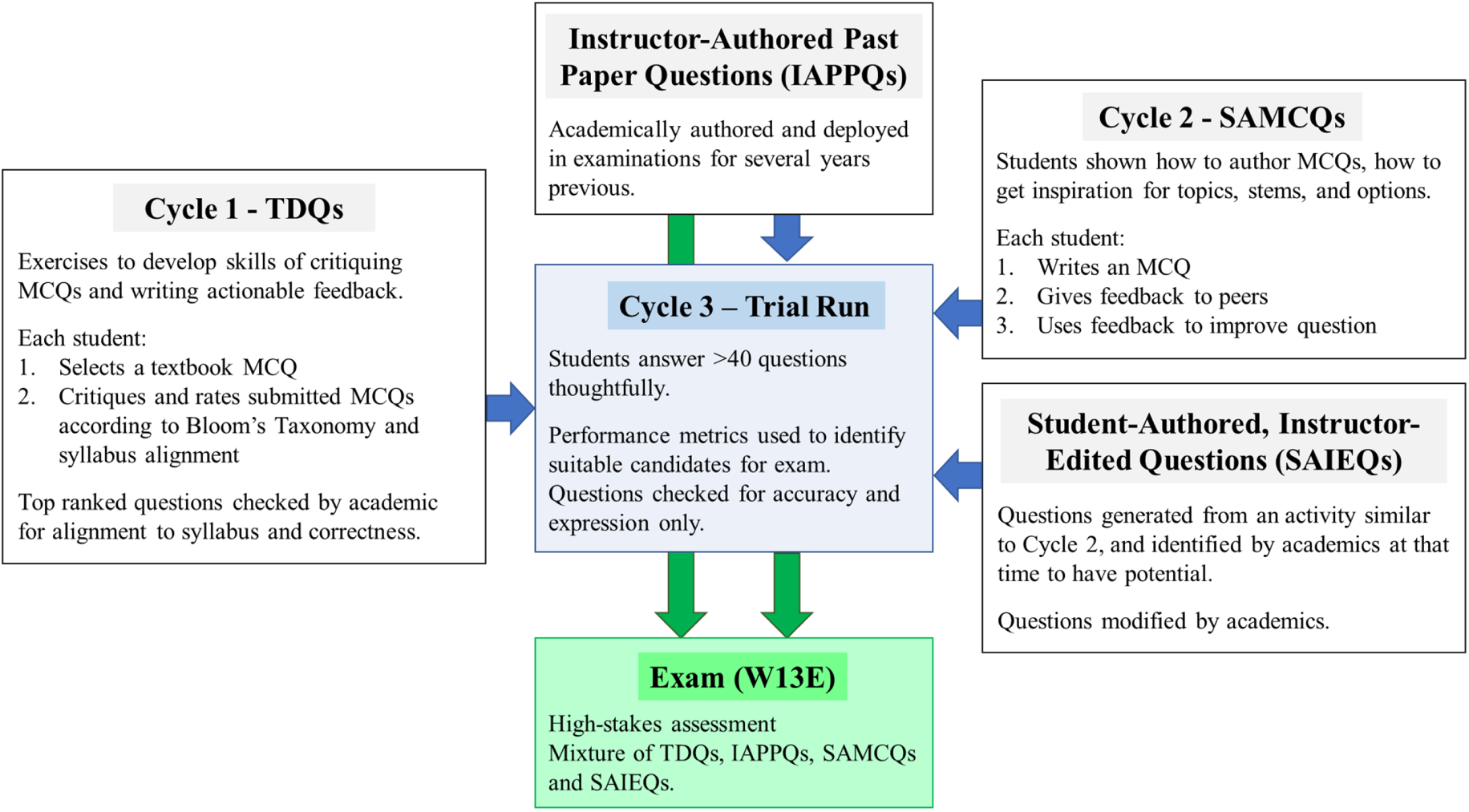
Summary of the process and the origin of the questions in the W13E.

### Details of Each Activity

#### Cycle 1: Reflecting on the quality of MCQs: learning to identify attributes and articulate opinion

In a tutorial setting, students were introduced to Bloom’s Taxonomy and its application to assessment items. Students discussed, in small teams of 5-6, the elements of seven sample MCQs, representing distinct styles and outcomes on the learning achievement spectrum.

The discussions were captured by a scribe on the collaborative whiteboard website Padlet (https://padlet.com/), and the sessions ended with a plenary discussion on those contributions. Students were also encouraged to submit their own reflections to PeerWise, both to gain familiarity with the platform, and to practice the language of Bloom’s Taxonomy. This activity generated several hundred pieces of feedback on each question and was effective in confirming that students were competent at recognising the key attributes of an MCQ, and were able to articulate their opinions.

The assignment for this tutorial was to source a question from a textbook or website on the topic assigned to their team (about 5 lecture slides; Table 1), and to post it for review. Each student had to contribute one question and review 10 other submissions.

**Table 1.**
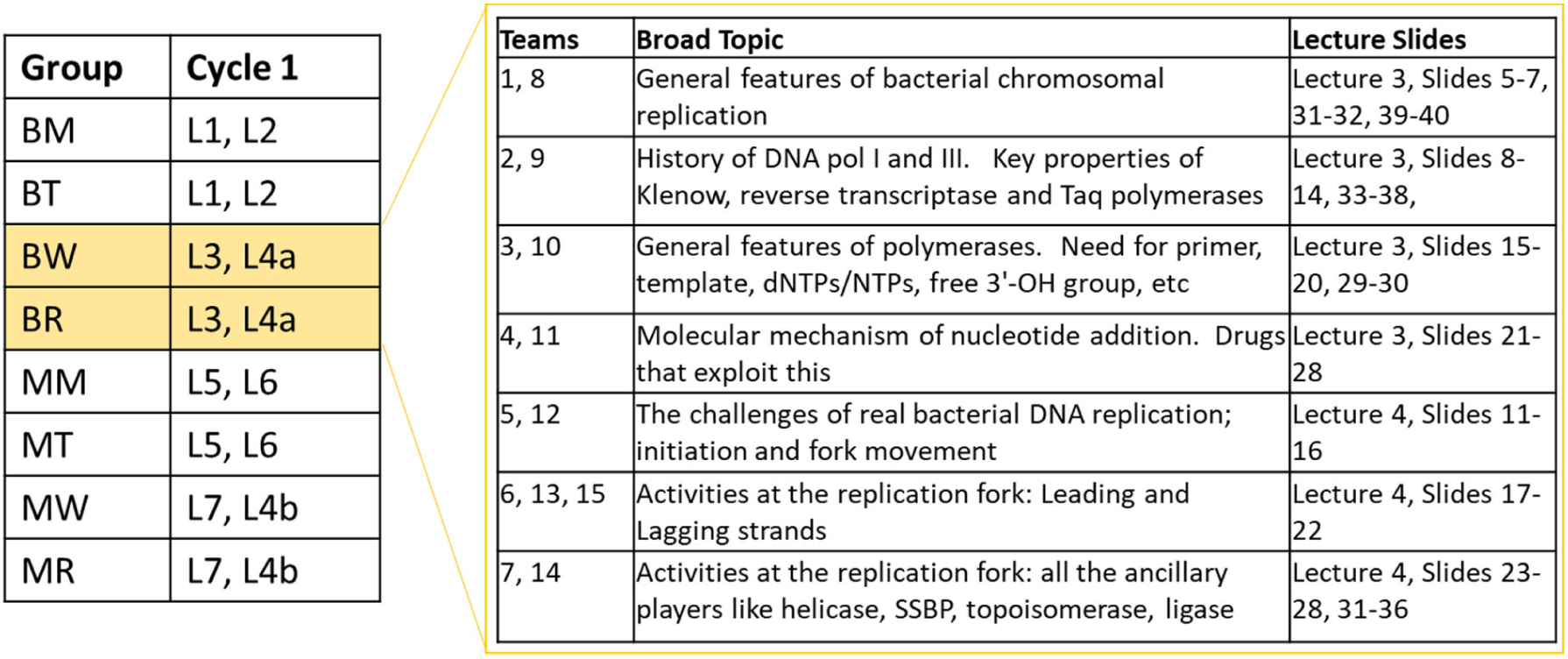
Setup of topic allocations in Cycles 1 and 2. There were eight tutorial groups of 70-80 students, which were further split into teams of about six students. For example, BW and BR were allocated the topics shown on the right. This was rotated for Cycle 2 (e.g., BW/BR worked on L5, L6 in Cycle 2).

Marks were awarded based on the quality of question review. It was also important that questions were submitted on time and appropriately tagged, but question quality was not marked. The differences in question quality was ideal for developing criticism and articulation skills, and critiques could be honest, as they were not authored by the students themselves. 667 questions were submitted, with 94% of these reviewed by more than 5 students, and 49% reviewed by more than 10 students. Students were also asked to judge each question based on Bloom’s Taxonomy using the standard PeerWise rating scale of 0-6 (0 for surface recall, to 6 for creativity/extrapolation), and difficulty scale of 0-2 (0 for easy, to 2 for hard).

After Cycle 1, students received feedback on their critiques and reminded that useful feedback has to be actionable and specific. They were given examples of feedback deemed ‘useful’ (thoughtful, specific, constructive, insightful, suggestions, expansive, articulate, comment on alignment to syllabus, reference to Bloom’s) and ‘non-useful’ (platitudes, perfunctory, general, repeated aspects of question).

About 40 questions from each of the eight groups were considered for deployment in future activities (i.e. the TDQs). To be eligible for selection, the question had to have at least 10 answers, a single most popular option, and a student rating on the Bloom’s Taxonomy level of at least two. These questions were then appraised by a subject matter expert (a teaching assistant in the course) to confirm correctness and alignment with the syllabus. This person could reject questions but were prohibited from changing the structure or wording beyond minor edits to improve clarity of expression.

#### Cycle 2: Question Authoring; Development and Refinement

This cycle aimed to equip students with the skills to harvest authentic peer misconceptions and profound insights on a specific topic, and integrate these into a novel MCQ. Students participated in a tutorial in which they explained concepts to each other, with members of the group challenging, querying, and extending these concepts to scope the boundaries of each other’s knowledge. Students then used this intelligence to each design a question on their allocated topic (Table 1).

As with Cycle 1, students were required to give feedback on questions submitted by their peers in line with Bloom’s Taxonomy. They were also asked to comment on question structure, and the extent to which it prompted reflection of the concept being tested. Student-authors could then use the comments to improve their submission. Only token marks were allocated to reward editing activity, but it was successful at encouraging revision of 78% of questions. No academic judgement of question quality was made, with marks primarily assigned to comment quality and action based on peer feedback.

#### Cycle 3: Determination of Performance Metrics of MCQs

All the questions from Cycle 2 were redeployed in new PeerWise courses to the classes that produced them (∼70 SAMCQs each). We also added ∼15 TDQs, 15 IAPPQs, and 15 SAIEQs to each of these pools. The SAIEQs were produced in a similar activity to Cycle 2, with the major difference being that they had been specifically assessed by academics, and had been tagged as being suitable for further development. Accordingly, these questions were revised by our team of academics to become SAIEQs.

Each of the ∼70 students in each class were tasked to answer at least 40 questions from their pool of around 110 questions. Although we were unable to control which questions were answered, >90% of the questions in each pool were answered at least 25 times. As this was a low-stakes task, it was also difficult to ensure that students thoughtfully considered each question. Since the purpose was to obtain accurate performance metrics, it was vital that students did not just choose easy questions or corrupt the data by answering flippantly. Therefore, to reflect an authentic exam setting, students were told that our scoring algorithms would reward genuine attempts, characterised by spending at least 1.5 min per question and submitting defensible quality and difficulty ratings. Students were not required to provide comments, nor were they assessed.

We judged this approach to be successful, as the average time spent by each student was 74.2 min ± 3.2 for the eight groups, which in total submitted over 24,900 answers. A custom dashboard was designed to easily view the outcomes of student answering activities. Data from students judged to not have taken the task seriously (generally <30 min, and undertaken close to the deadline) were omitted from analyses of question performance metrics. Within each of the eight groups, and based on the proportion of questions answered correctly, students were divided into tertiles for the computation of question performance metrics.

The main metrics calculated for each question were difficulty (% correct) and discrimination index (DI; the proportion of the bottom tertile that answered correctly subtracted from the proportion of the top tertile that answered correctly).

#### Use of Intelligence from Cycle 3

Questions with a DI >0.2 and a difficulty of ∼60% were considered candidates for inclusion in the high-stakes assessment. This filtered the pool of 702 questions to ∼200, including all the IAPPQs, which were automatically included. The performance of the distractors in each of these questions was classified according to two easily implemented heuristics; a) identification of obvious ambiguities, as shown by students selecting one or two major options, and b) reflection on the utility of each option, with particular consideration given to the identification of options that were not being picked at all. Although in this iteration this task was performed manually, these attributes can be calculated either for automation or to assist the decision-making process.

Questions with minimal ambiguity and at least three selected distractors were then quality checked by an academic. Only minimal editing was performed; sufficient to confirm that the question conformed to baseline standards, aligned to the syllabus, interpretable, and had a single, genuinely correct, option.

### Preparation of High-Stakes Assessment

Four sets of 42 questions each were prepared for four different versions of the W13E. Each of these assessments contributed 10% to their final marks, and were run in a timed (1 hour), online format. Although, from an assessment point of view, we wished to include only questions with ideal DI/difficulty metrics from Cycle 3, we compromised to ensure that each paper had approximately an equal blend of questions from the four sources. The four exams were deployed throughout the week, allowing students to choose which day to sit the task. For fairness, we ensured consistency among question pools in overall difficulty, DIs, expected response time, and coverage of learning outcomes.

### Determination of Question Performance

In addition to calculating the difficulty and discrimination factors by traditional methods, we employed two other approaches.

Item Response Theory (IRT) was used to generate a graphical depiction (Item Characteristic Curves) of how the probability of success within a question was related to student ability. It allowed for a more granular and dynamic appraisal of the relationship of these factors than the traditional DI. To generate the curves for each question, the score obtained by each student was processed in the statistical analysis package, R, according to the workflow designed by Xie, Davidson and Ko (2019).

Briefly, W13E data was organised in a spreadsheet with each row representing a student and each column a question, so that each cell contains a student’s response (1 = correct, 0 = incorrect) to a specific question. This table was submitted to the “ltm” package for R, which displays the probability that students of particular ability would get each question correct. Examples of outputs typical of high- or low-performing questions are shown in the Results.

Distractor Frequency Analysis (DFA) is an in-house developed method for describing the likelihood that cohorts will select particular options within an MCQ. The class was divided into six groups based on overall exam or MCQ component mark, and the percentage of students in each sextile that choose each option was calculated and presented in an easy-to-interpret interface. Examples of outputs typical of high- and low-performing questions are shown in the Results.

## Results

### Monitoring Student Contributions

A key component of our strategy to train students to become effective MCQ authors was to encourage them to provide timely, actionable feedback, and was therefore our focus in Cycles 1 and 2. PeerWise provides a report that collates all the contributions (questions, comments and replies) from one student on one page. It takes about 30 seconds to scan the comments and confirm their usefulness or otherwise (Figure 2).

**Figure 2.**
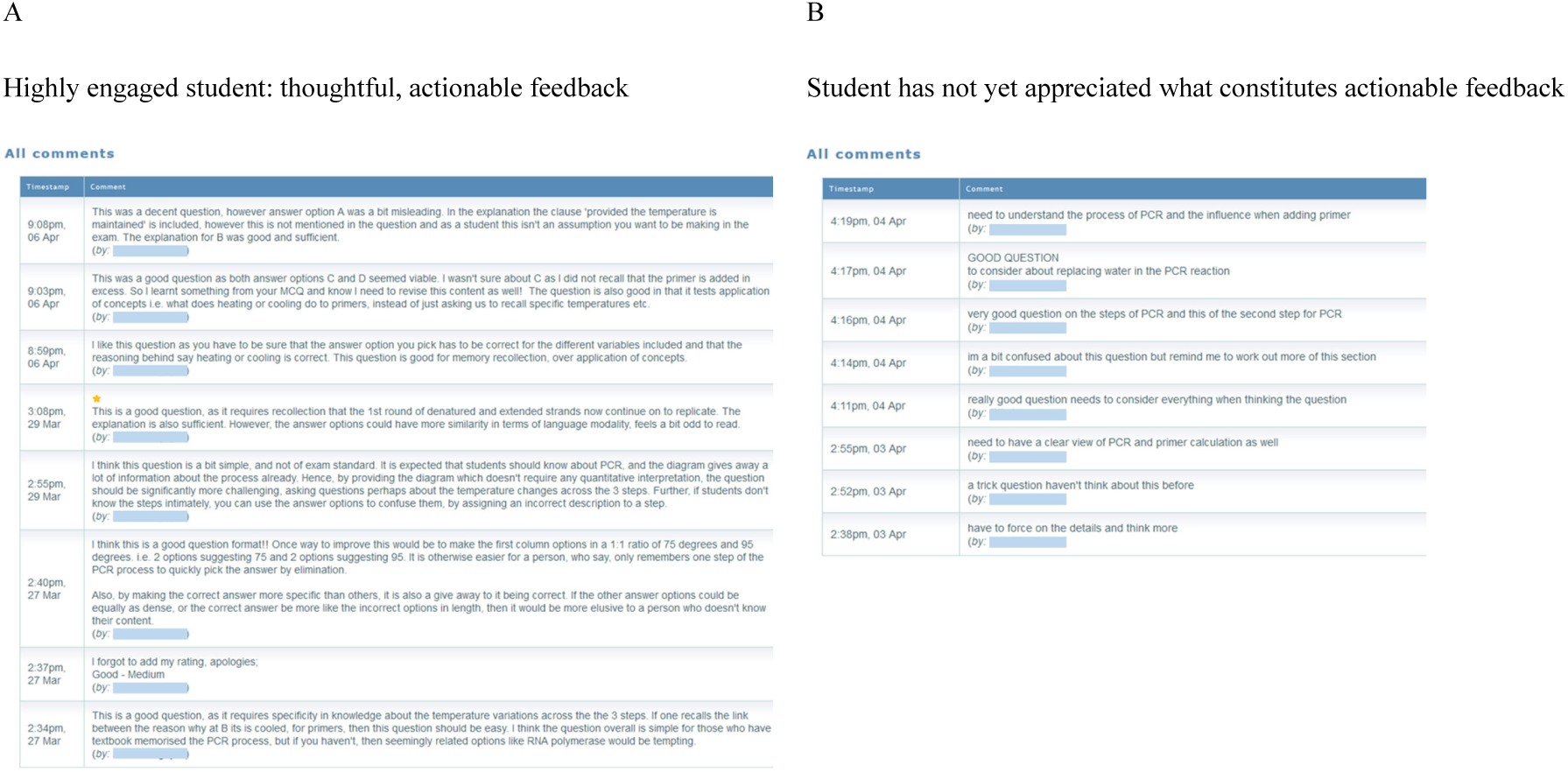
Comments of an engaged and not so engaged student to the same questions in Cycle 2. The feedback from Student A can be used to improve the question. Moreover, they are more likely to reflect more on the attributes of their own question. Outputs like this were valuable in showing students how they should develop their articulation skills.

Success of the broad question performance trawl in Cycle 3 depended on students approaching this largely formative task with sincerity, answering each question to the best of their ability. From experience, students generally take a task seriously if they trust that their efforts will be rewarded. Therefore, we developed a dashboard (Figure 3) using granular time-stamped activity data (available on request from PeerWise), to convince students that we could see their approach to the questions. This strategy proved successful as the vast majority (>87%) of students completed the task with appropriate diligence, judged only from inspection of the start and end times of answering sessions. Such students spent 1-2 hours answering the 40 questions, and, reassuringly, 12% of students spent less than 1 min per question. Only 5% were judged to have not performed the task thoughtfully, as evidenced by them spending less than 30 seconds per question, selecting mainly ‘easy’ questions, and still answering the majority incorrectly.

**Figure 3.**
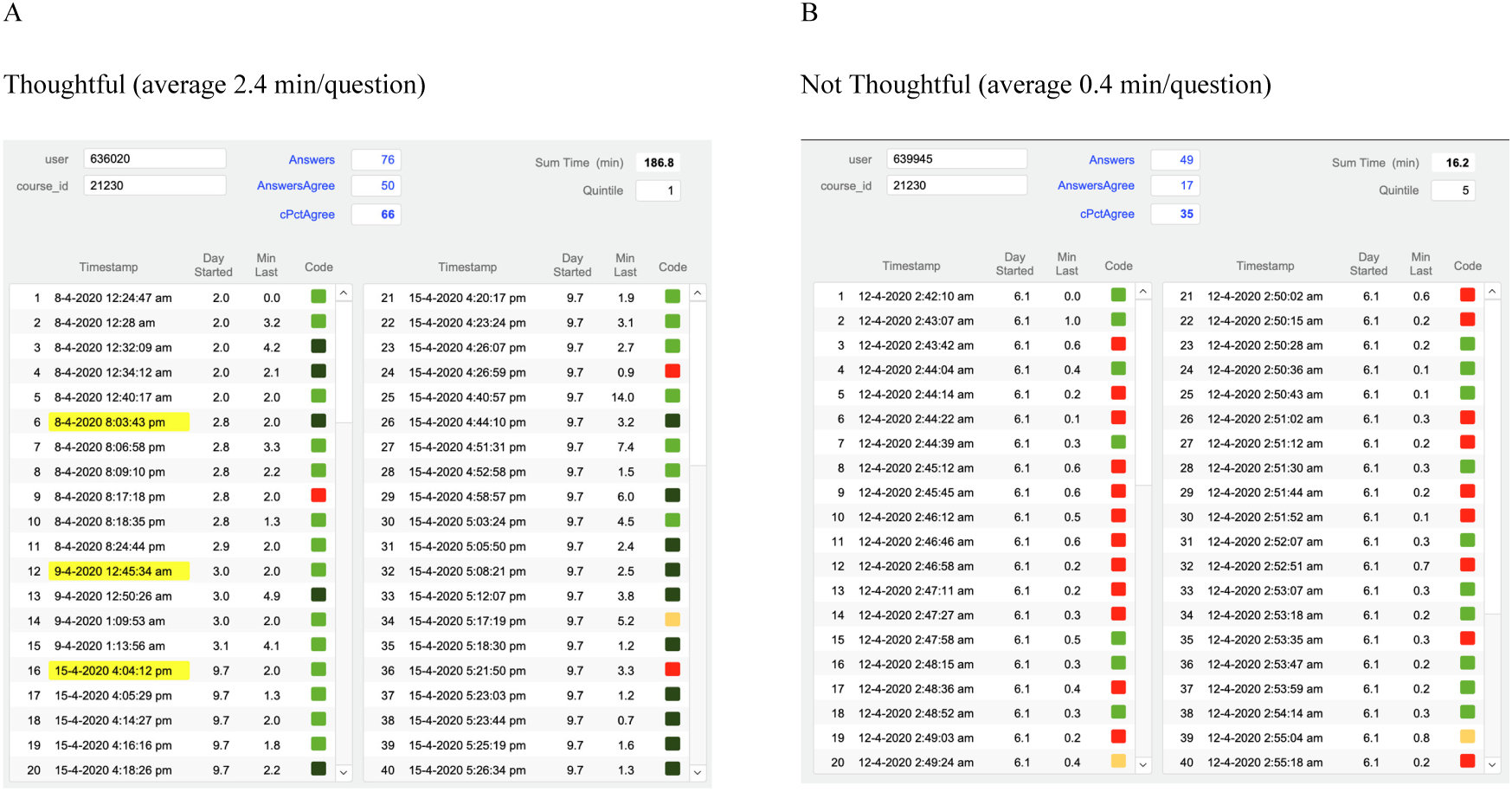
Dashboard developed to monitor student engagement in Cycle 3. Answering patterns are shown, including successful attempts on difficult questions (dark green), unsuccessful attempts on easy questions (red), and commencement of any sessions following the first one (yellow highlights). Data from students such as Student B were excluded from analyses of question performance.

### Overall Exam Information

Over 600 students sat one of the four W13E, with nearly 55% of them taking the final, Thursday paper. Despite each exam consisting of largely different questions, with only 2-3 questions being redeployed on other days, each paper was of approximately equal difficulty. Both the average and lowest exam scores were reduced for the Thursday version (Table 2), but the cohort of students who took this exam were slightly weaker, as indicated by their performance in the formal end-of-semester examination, assessment of the remaining lecture content that contributes 30% to their final mark (Figure 4). We did not include questions from the Monday exam in further analyses, due to the low number of students.

**Table 2.**
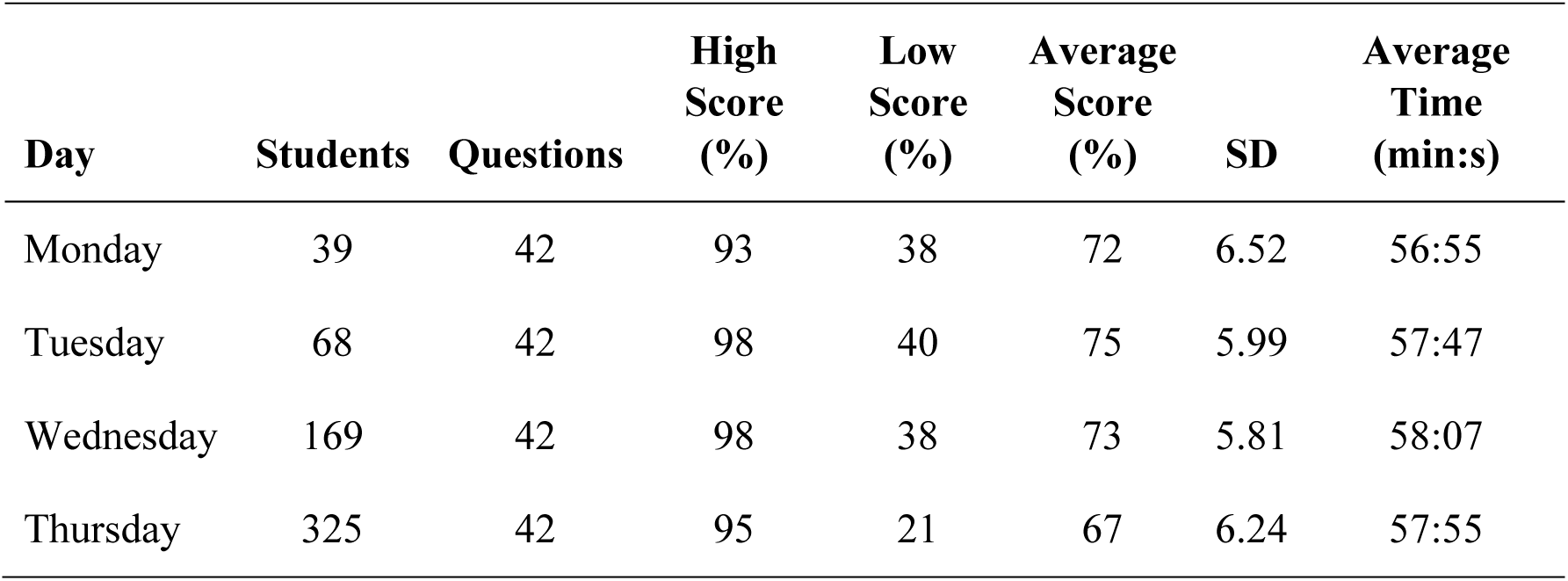
Statistics of the four W13E.

**Figure 4.**
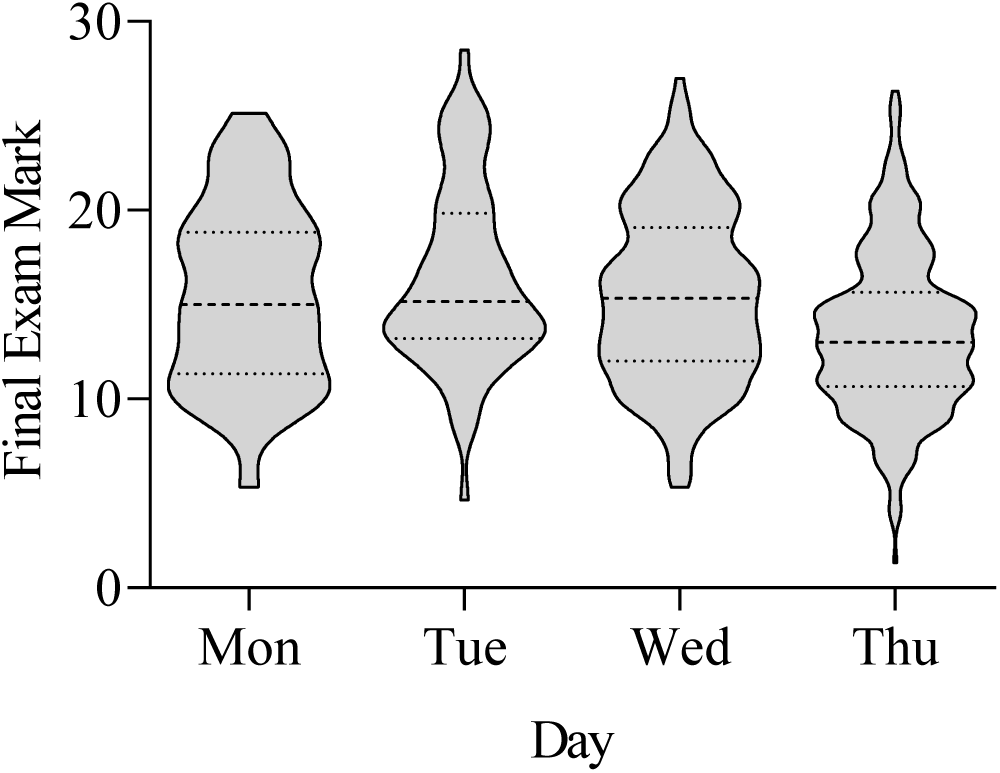
Performance of students in the Final Exam. The marks of the end-of-semester exam grouped by the day on which the students sat the W13E. The dashed and dotted lines represent the median and quartiles, respectively.

There was a strong linear relationship between the W13E (containing >50% student-authored questions) and the marks of the Final Exam (all academic written questions) with the correlation being particularly strong for the higher-performing students (Figure 5).

**Figure 5.**
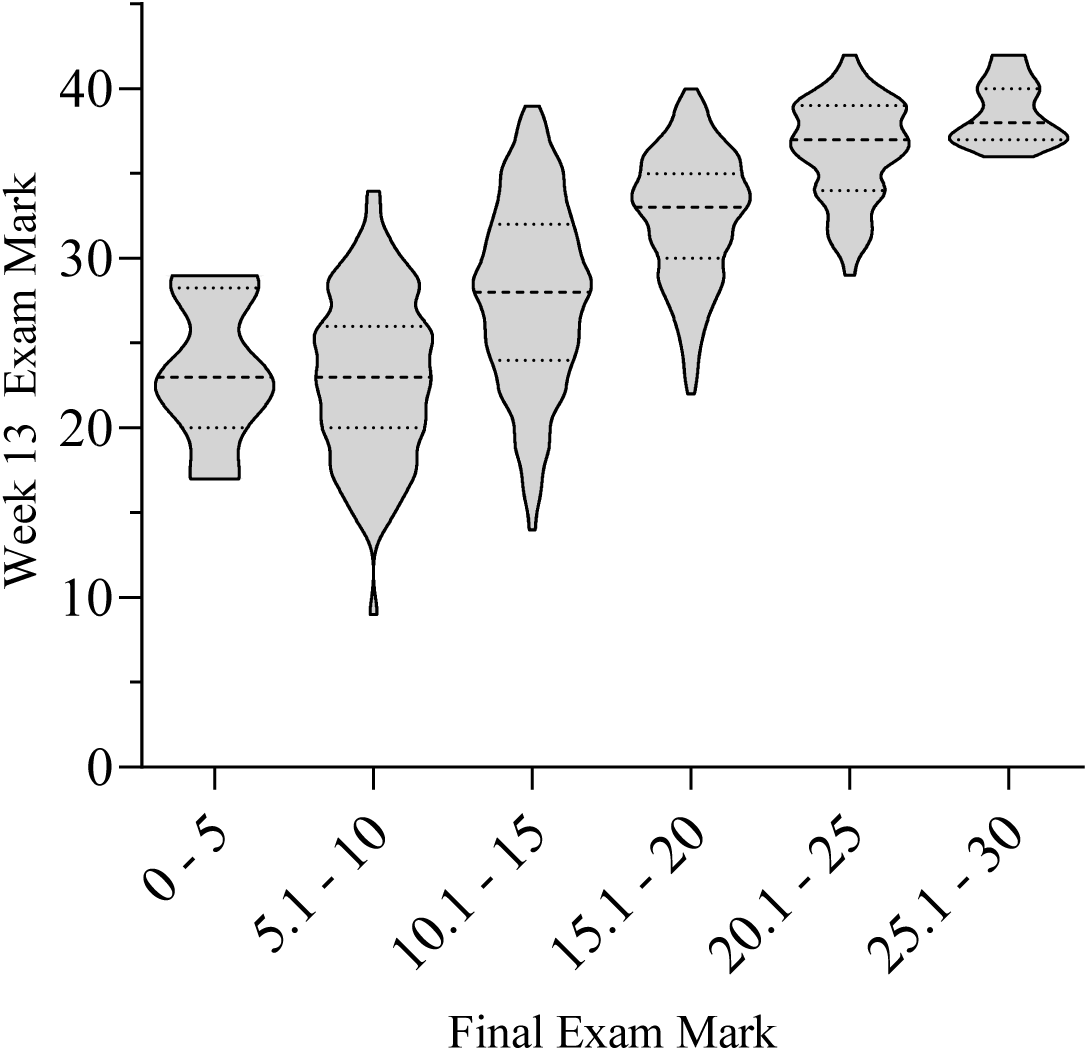
Comparison of student performance in the W13E and end-of-semester exam. Marks from the W13E were plotted for each student, grouped by their performance in the end-of-semester exam. The dashed and dotted lines represent the median and quartiles, respectively.

### Question Performance Analysis and Classification

The performance of each of the 126 questions from the Tuesday, Wednesday and Thursday exams was assessed according to three separate analytical techniques; Classical, IRT, and Distractor Frequency Analysis (DFA). In the IRT, the ability of the students is plotted against the probability that they will answer the question correctly. An ability of 0 represents students performing at an average standard, and 4 and −4 representing the highest and lowest achieving students across the entire exam.

In the example (Table 3), the probability that a student of average ability will get this question correct is over 0.6, and over 95% of high-achieving students and less than 5% of students at the bottom end are getting it correct. The strong performance of this question is supported by the Classical analysis, which shows that 64% of the class chose the correct option (B), and that the DI for this option (and, therefore the question as a whole) was 0.33, i.e., the proportion of students choosing option B was much higher in the stronger students than in the bottom sextile. Note that the false option, A, shows negative DI, which is desirable.

**Table 3.**
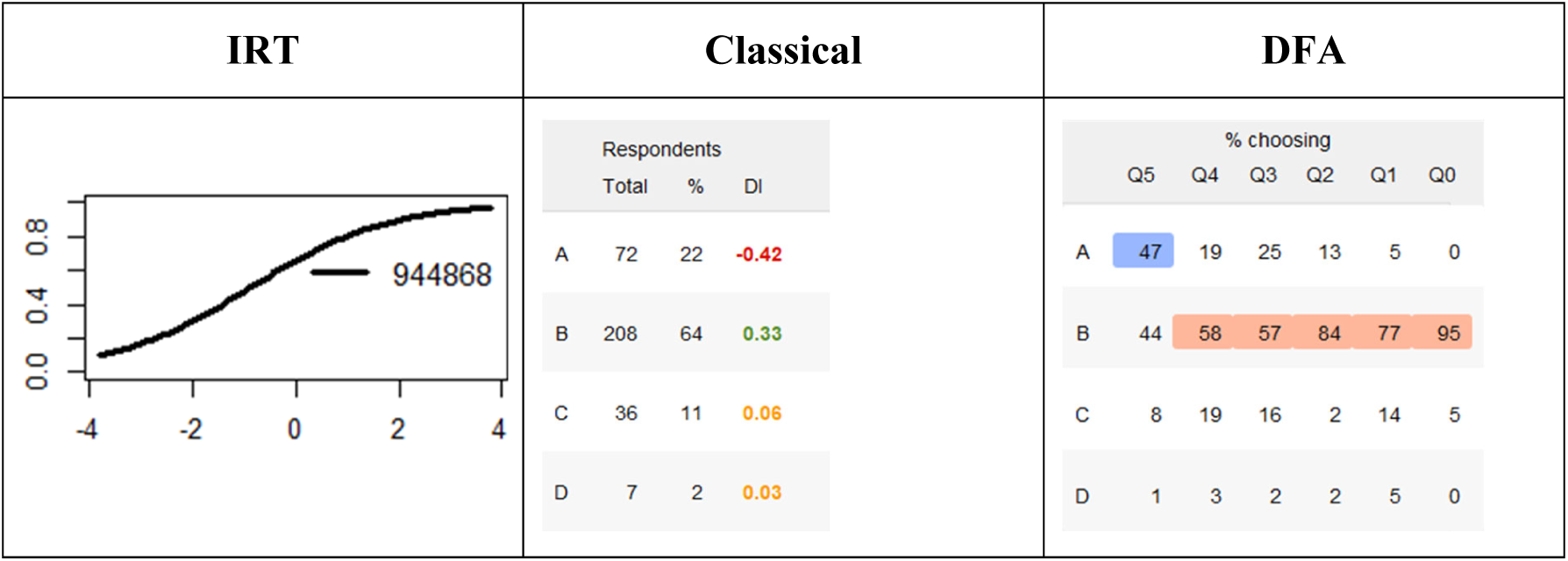
Performance metrics for ‘Good – Standard’ question. The x-axis of the IRT analysis represents ability, and the y-axis represents probability that they answer correctly. This appeared in the Thursday paper, in a random position for each student. The number 944868 is the unique ID of the question allocated within the online delivery platform.

DFA expands on the classical approach by showing the proportion of each sextile that choose a particular distractor. Q0 is the top sextile and Q5 is the bottom. Correct choices selected by more than 50% of the sextile are highlighted in pink and wrong choices selected by more than 25% are in blue. This complements the other analyses by providing intelligence on the likelihood that particular cohorts will select specific options, and if the options appear ambiguous to particular groups of students, which may or may not be desirable.

Using performance metrics, we classified each question into five main categories: Good – Barrier (deliberately easy questions that all moderately engaged students should answer correctly), Good – Standard (discriminating effectively between passing and higher achieving students), Good – Difficult (discriminating between students at the top of the class), Poor – Noise (answered effectively randomly by students of all abilities), Poor – Negative (actively penalises the higher achieving students) (Table 4). Categorisation of each question was agreed on by each researcher with little ambiguity.

**Table 4.**
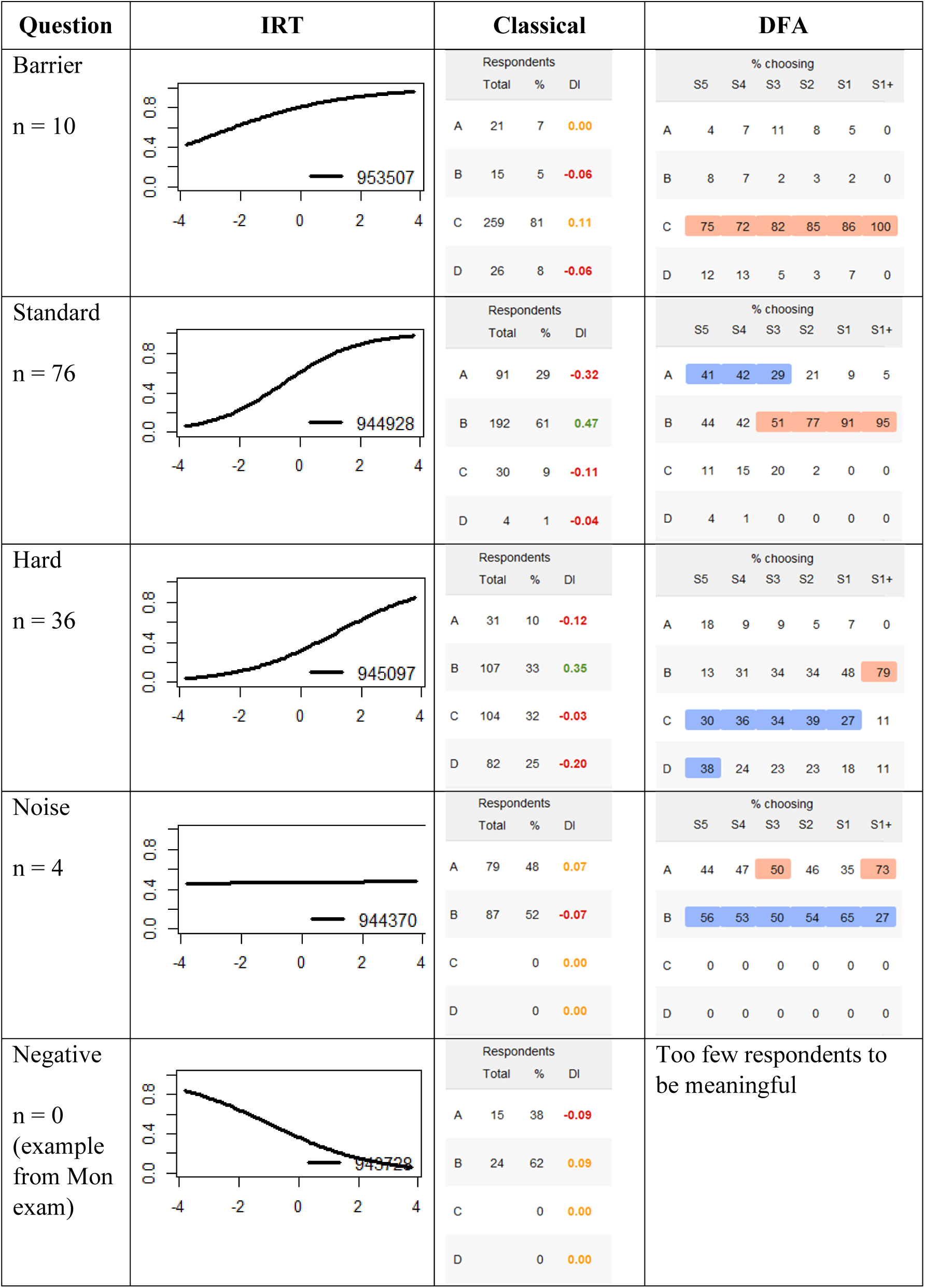
Summary of the types of questions that appeared in the exam. The x-axis of the IRT analysis represents ability, and the y-axis represents probability that they answer correctly.

For very easy questions, the IRT curve is flat and consistently high, with students at both top and bottom sextiles answering this question correctly, and with none of the distractors being attractive to any particular cohort. It is arguable as to whether a question of this nature should be classified as ‘good’ or ‘bad’. While it does not discriminate between high- and low-achieving cohorts of students, if pitched to confirm achievement of a core learning outcome, it is a worthy measure of student achievement. Academics are increasingly encouraged to include barrier questions such as this, but only 10 of the 126 questions in our exams were so classified.

The majority of questions (76/126) in our exams were classified as a good standard question, which discriminated between passing and failing students. The IRT curve shows that students in the higher ranges of ability all answer this question correctly, and even those of medium ability get it correct more than 50% of the time, whereas very poor students do not get it correct. This is supported by the traditional metrics, which show a correctness/difficulty of 61% and a DI of 0.47. Further reinforcement of the classification of this question comes from the DFA, which confirms that over 90% of the top sextile and around 40% of the fourth sextile (representing the passing students) answered it correctly. However, the extra intelligence obtained from DFA is that Option A was particularly attractive to weaker students and Option D was a very weak distractor for all cohorts.

Questions were classified as hard if they discriminated between students towards the top of the class. The IRT curve is shifted to the right, the overall correctness is reduced to 30% and the DI is >0.3. In addition, the DFA confirms that 80% of the top sextile got the question correct, with every other sextile favouring Option C except the weakest students who opted more frequently for Option D.

One indication of a poor question is a flat IRT curve in which the question is answered effectively in a random manner across all abilities. Indeed, DFA and Classical analyses confirm that students were reduced to taking a 50:50 bet between two obviously ambiguous options. A question like this generates noise in the student scores but does not have an overtly negative effect on either cohort.

The worst type of question is revealed in the last row of the table. The IRT plot has a negative slope, the DI is negative, and Classical analysis shows that students were choosing between two options. However, the better the student, the more likely they were to select the option that had been classified as incorrect. None of the 126 questions used in our analyses were classified in this way.

### Analysis by Traditional Metrics

Based on traditional metrics, the SAMCQs and SAIEQs were at least as good as academic-authored questions, whether they were past paper questions or those sourced from the textbook or web (Figure 6A). Indeed, contrary to the latter two groups of questions, the DIs of all SAIEQs were positive. Questions from the textbook/web contained several questions with low or negative DIs, and the greatest spread in the DI range was seen in the past paper questions.

**Figure 6.**
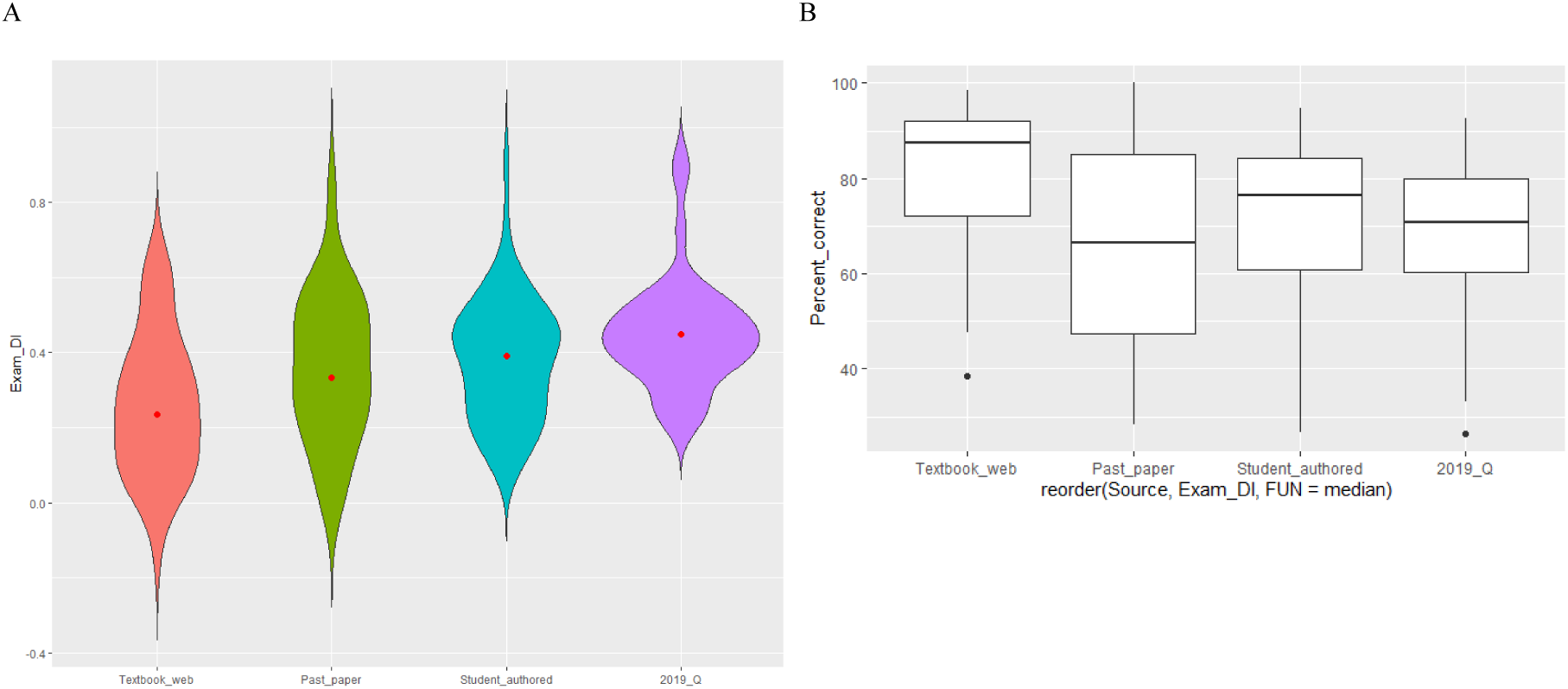
Comparison of traditional metrics of questions from the four sources. The sources from left to right: TDQ, IAPPQ, SAMCQ, SAIEQ, and their (A) discrimination indices and (B) difficulty.

Textbook and web-sourced questions were also overall easier than student-authored and past paper questions (Figure 6B). As with DIs, the past paper questions also had a greater range of difficulties than the student-authored questions. Student-authored questions with and without contribution by an academic exhibited a smaller range and were of similar difficulty to the harder past paper questions.

### Internal Calibration of Past Paper Questions

The performance of the IAPPQs was particularly important to define as this serves as an internal control, allowing us to determine if a) the class were of a similar standard to previous cohorts, and b) we were getting the students to engage with the material in a similar way despite some changes in lecturing staff. This was particularly important, since, the 2019 questions were part of a larger, traditionally administered examination that covered the entire syllabus including both molecular biology and metabolism concepts (15 lectures of each). In contrast, the 2020 questions were part of a smaller, shorter, online MCQ exam that covered only half the molecular biology material (7 lectures). Another difference between the two years was that the 2020 students had the option to choose which of the four exams they sat.

IAPPQ8 was typical of most of the easier IAPPQs, with students towards the lower end of the class getting this right more than 50% of the time and with identical DIs and derivative curves (Table 5). IAPPQ11 performed similarly between the 2020 and 2019 exams, even though it was a harder question (as illustrated by the peak in the derivative curves at higher ability students), and despite it being run on two separate exams in 2020. IAPPQ6 behaved slightly differently between the two years, with the 2020 deployment being a better discriminator largely because the less able students answered it more poorly. Overall, 12 IAPPQs were deployed in both 2019 and 2020 and, in every case, the performance of each was consistent with the examples shown in the table.

**Table 5.**
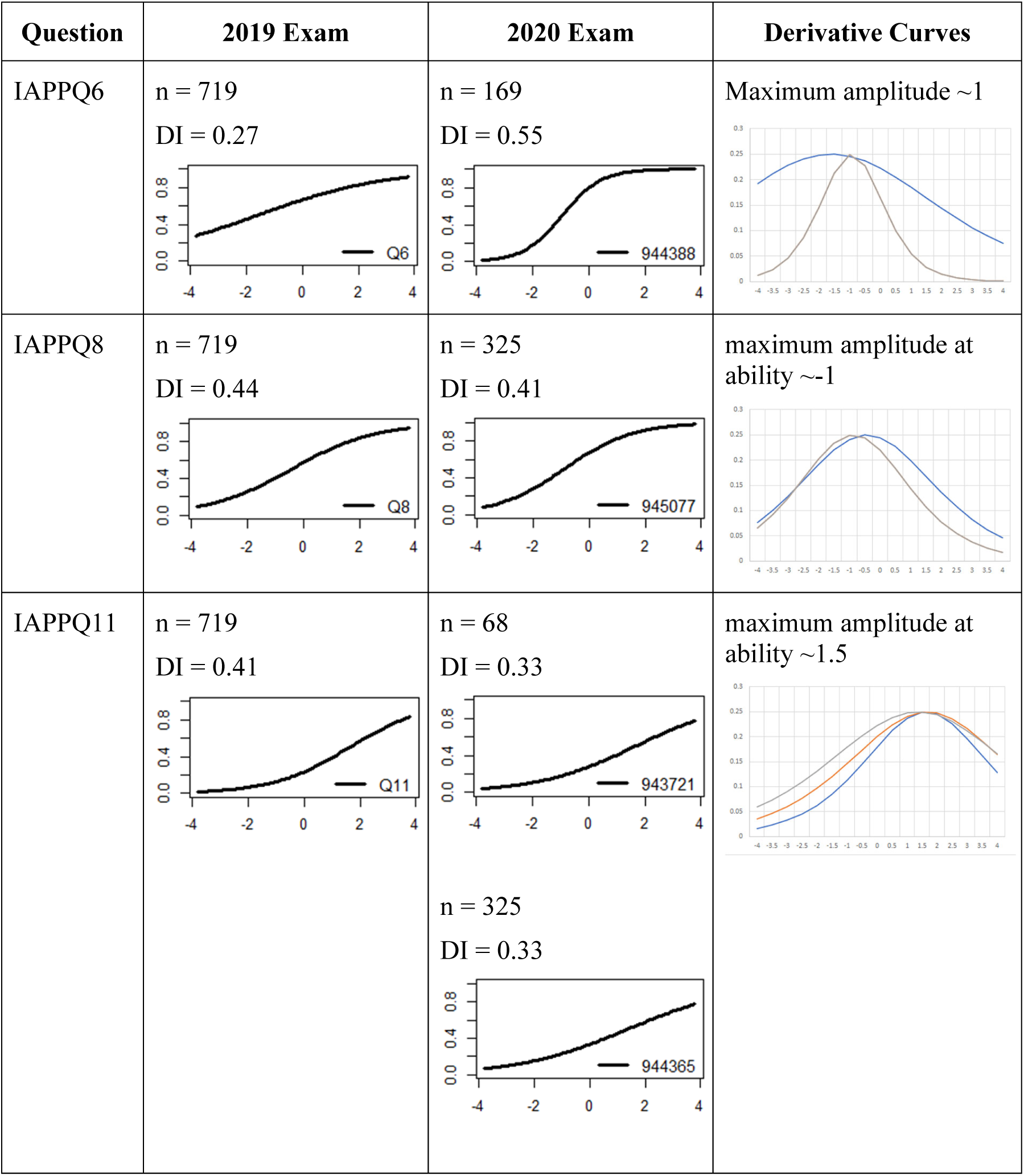
Comparison of performance of past paper questions. The x-axis of the IRT analysis represents ability, and the y-axis represents probability that they answer correctly.

The peaks of the derivative curves are the inflexion points of each IRT curve, i.e., the ability level that gives a 50% probability of getting the question right. The spread indicates the extent to which the question discriminates students around this point. The curves were obtained by plotting the first derivative of the IRT plot equation against the ability level (−4 to 4). Sharp peaks are items with strong discrimination between the top and bottom of the cohort, while broader peaks show less discrimination.

Using the 2-Parameter Logistic Model, the probability of student *i* correctly answering question *j* is given by:

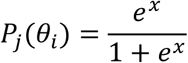

where *x* = *a*_*j*_(*θ*_*i*_ – *b*_*j*_), *a* is the discrimination of the question, *b* is the difficulty of the question, and *θ* is the ability of the student. Thus, a student has a 0.5 probability of answering a question correctly if student ability equals the difficulty of the question.

The first derivative of the equation above is:

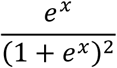

Using the difficulty and discrimination coefficients calculated by the R model program, it was thus possible to calculate and plot ability levels between −4 and 4 using this equation to produce the first derivative plots.

### Analysis by IRT and DFA classification

All of the student-authored questions that had been edited by an academic were classified as good, and contained both easy and hard questions. The past paper questions also performed similarly, with the addition of barrier questions. Out of the four questions that were categorised as poor, two of these questions were textbook and web-sourced questions, and two were student-authored questions that had only been minimally edited (Table 6).

**Table 6.** Quality of questions from the four sources.

|  | **TDQ** | **IAPPQ** | **SAMCQ** | **SAEIQ** |
| --- | --- | --- | --- | --- |
| **Good** |  |  |  |  |
| - Easy | 19 (63%) | 11 (55%) | 27 (59%) | 19 (63%) |
| - Hard | 4 (13%) | 7 (35%) | 14 (30%) | 11 (37%) |
| - Barrier | 5 (17%) | 2 (10%) | 3 (7%) | 0 (0%) |
| **Poor** | 2 (7%) | 0 (0%) | 2 (4%) | 0 (0%) |
| **Total** | 30 | 20 | 46 | 30 |

Indeed, out of the 126 questions analysed, there were only three questions with particularly low DIs; two of these questions were sourced from the textbook/web, with the other being a student-authored question that had not been edited by an academic.

### Comparison of question performance in low- vs high-stakes tests

The low-stakes formative activity in Cycle 3 was an important part of the process of selecting suitable questions for the high-stakes summative W13E. Presumably, students were taking more care when completing the summative assessment than the formative activity. To determine the extent to which the metrics could predict the suitability of questions in a real exam, we compared DIs (computed on the basis of tertiles) and difficulties of the questions in the two exams (Figure 7). There was broad agreement in question performance between the exams, particularly in the DI range 0.2-0.6. However, there was a greater spread at very high and low exam DI levels. This is likely because the Cycle 3 DI values are based on fewer responses and a less precise scale of esteem, and are therefore less reliable. As expected, a greater proportion of students answered correctly in the exam than in the low-stakes assessment.

**Figure 7.**
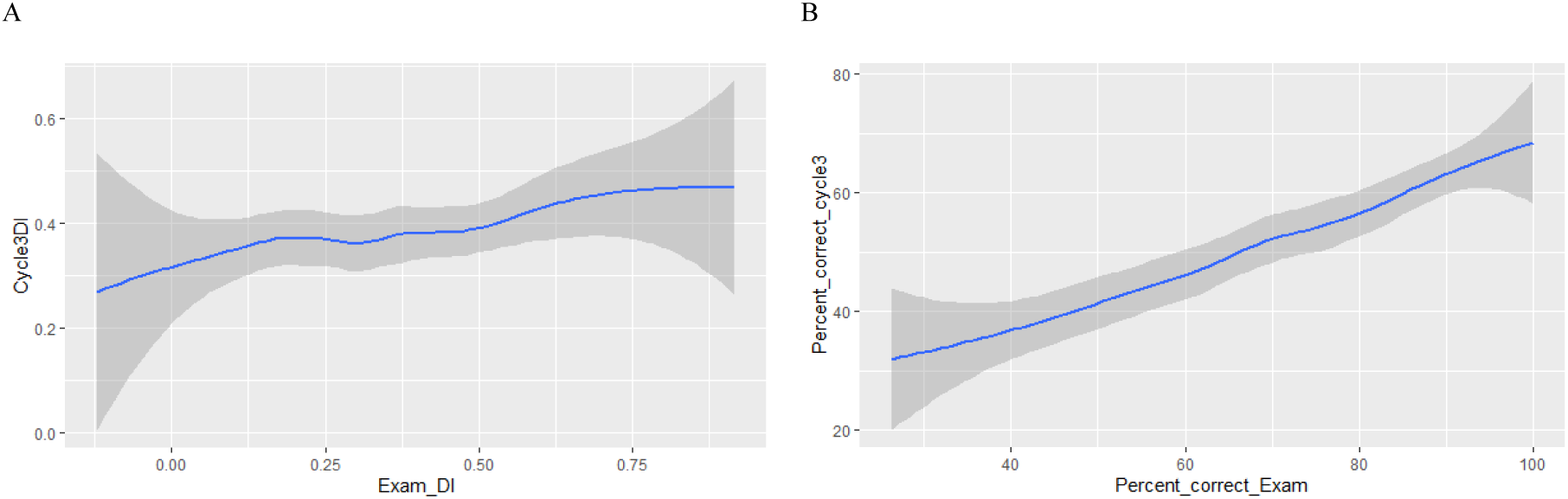
Comparison of question performance in a low-stakes and a high-stakes assessment. The (A) discrimination indices and (B) difficulty of the W13E were plotted against metrics calculated from Cycle 3.

## Discussion

In this study, we describe a scalable approach for training students to author high quality MCQs, and we compare two approaches for selecting student-authored questions to appear on summative assessments. We found that questions selected by either approach performed as effectively as academic-authored past paper and textbook-sourced MCQs.

Several studies have reported crowd-sourcing questions from students, not only to build up revision banks, but also as a form of revision itself, since the process of authoring and critiquing questions has been shown to raise student performance. However, because high-stakes summative assessments must be error-free, student-generated questions are rarely used directly for this purpose (Schullo-Feulner et al., 2014). This seems to be the case even when the questions have been subject to quite stringent academic review. For example, Harris et al. describe an activity where student-authored questions were sequentially reviewed by their peers and experts for factual accuracy and distractor quality, with approved questions published for test practice (Harris et al., 2015). However, the questions were not deployed in high-stakes summative exams. Amini et al. went a step further, combining approved student-authored questions with teacher-authored questions in a medical imaging exam for radiology students (Amini et al., 2020). Analysis of the exam responses revealed that student-authored questions were easier, had significantly more non-functional distractors, and were more likely to rely on recall skills compared to teacher-constructed questions. However, they did not report any explicit training or coaching of students in question authoring, and unlike us, did not actively select student-authored questions with a high discrimination index.

### Two approaches for selecting student-authored questions

We examined two distinct approaches for selecting student-authored questions for deployment on a high-stakes summative exam:

1. identifying candidates by trialling the questions in low-stakes quizzes, followed by light editing to ensure compliance with learning outcomes and basic delivery standards.
2. identifying potential questions using manual academic judgement followed by intense editing by subject matter experts.

Both methods delivered suitable questions but each comes with quite different overheads in terms of workload and practicability. Filtering and candidate selection using a list of automatically generated performance metrics from low-stakes quizzes is much easier and faster than academic selection involving manual assessment of the stem, options, structure, logic and expression of individual questions.

A key advantage of the low-stakes trial approach was that it provided a reasonable indication of how the questions were likely to perform in the summative exam, something that is difficult to predict from untrialled questions, even for experienced academics. In addition, the selection criteria can be varied to deliver barrier-style (high easiness) or higher-level MCQs as required. Additionally, selection is not biased in the structure or sentiment of the question. In contrast, each academic will have a bias for/against certain MCQ structures and formats. Questions written by students may be dissonant to an academic’s normal style, and it can be unsettling to run an assessment using these questions. The increased richness in the range of question styles produced by students provides an opportunity to new ways of thinking and is strongly student-centred, by definition arising from their own stylistic preferences. When faced with the manual selection approach, we noted that as academics, we tended to choose questions that show creativity, insight, extrapolation, and effort. This leads to the selection of items that are at the higher levels of Bloom’s Taxonomy. However, more complex questions require more work to ensure it is not ambiguous and that each option functions as a suitable distractor. The reward is potentially high because there is a strong chance that the process will produce questions that are innovative and discriminatory, and are also strongly aligned to the values, objectives, and style of the individual academic, but the time investment is substantial.

A caveat to the low-stakes trial approach is that the metrics can be deceptive. Poor questions can have a high DI if high-achieving students are gravitating to the least incorrect option. Conversely, potentially excellent questions can have a poor (or even negative) DI because strong students are choosing incorrect options that are unattractive to average ability students. For this reason, selection solely based on metrics is not recommended, but it is an efficient initial filter. Similarly, academic judgment alone is not always a reliable predictor of question performance. For example, SAIEQs that did not perform well in Cycle 3 were not included in the W13E. It is likely that, with the benefit of reflection on the performance metrics, some extra revision of the questions would have provided sufficient remediation.

Regardless of the question selection process, using published student-generated questions on high-stakes examinations introduces a concern that students who either authored or answered the questions prior to the exam may have an unfair advantage. There is evidence that students perform significantly better on subsequent exam questions if they have previously authored or answered questions that merely target the same topics (Kelley et al., 2019). Similar research, which addresses a potential bias relating to topic-selection, has shown that these results are robust even when students are randomly assigned topics on which to author questions (Denny et al., 2017). This concern is partially mitigated in our study as there was a very large question pool and all students had exposure to the entire repository before selection. One potential solution to eliminate any direct advantage would be to stagger the authoring and filtering tasks between academic years or cohorts of students.

### Sourcing Questions from Textbooks

Intuitively, textbook-derived questions (TDQs) would seem to provide the most efficient option, since each question has, presumably, been crafted to reflect fundamental concepts and may even have been tested in genuine assessments. However, even though we chose our 30 TDQs from a pool of some 600 candidates, guided by data from student ratings, and tempered with academic judgment, with reference to our specific learning outcomes, these performed no better than any other type of question. Our perception that TDQs are mainly useful for assessing generic facts was supported by the fact that 80% were classified as ‘easy’ or ‘barrier’, with just 13% being ‘hard’. Student ratings in Cycle 1 rarely rated the TDQs above the most basic levels of Bloom’s Taxonomy. Another issue with TDQs is that most academics present their subject matter with their own distinctive emphasis, and so it is often difficult to make ‘generic’ TDQs feel relevant to a bespoke syllabus.

### Sourcing Questions from Academics

Whilst it was not practical to obtain an objective measure of the workload associated with construction of IAPPQs, because these already existed, by definition, we are confident that our introductory assertion – that good quality, field-tested questions are precious – would resonate with most academics.

It would be inappropriate to suggest that all academic-authored questions are as strong as our IAPPQs, which have been constructed by our most student-centred colleagues and fine-tuned over many years of deployment. Indeed, it would not be uncommon for course coordinators to feel some disappointment with questions received from academic colleagues. MCQs from less student-engaged academics can often be hastily constructed, and without sufficient insight in the abilities of the student cohort. It is our perception that some research-focused academics favour writing questions that are pitched towards the high-achieving students, and many do not review the performance of their questions, nor refine them from year to year. Of course, novel academic-created MCQs all suffer from not being field tested or subjected to the same cyclical process of review, discussion and editing that occurs in a tool like PeerWise.

We would argue that academics could also benefit from classes on the strategic approach to MCQ authoring. It is sometimes hard for us, as expert practitioners, to see the topic through the lens of a developing learner. Even when given a very specific concept on which to design an MCQ, it can be difficult to know what facet of that topic should be assessed and the academic level at which the question should be pitched. Conversely, when our understanding is developing, it is the moments of enlightenment that define our advancing mastery of the topic, and it is these that form the basis of the most meaningful correct options in MCQs.

### Training and Associated Workload

We adopted PeerWise in the current study, as it is free to use, has excellent in-built reports on student activity, provides low-level access to data for the construction of bespoke dashboards, and has been a fixture in our course for several years. The benefits to student academic performance result from all aspects of engaging with PeerWise: answering questions, articulating criticism of peers’ questions, authoring questions, and editing questions in light of peer feedback (Doyle & Buckley, 2020; Hancock et al., 2018; Hardy et al., 2014; Hudson, Jarstfer & Persky, 2018; Kay, Hardy & Galloway, 2018; Kay, Hardy & Galloway, 2020; Walsh et al., 2018). Engaging in discussions that reveal insights and misconceptions, and the related reflective and critical processes involved, are likely to be valuable components of the learning experience. In our study, we put an emphasis on training students to create and criticise MCQs through scaffolded tutorials and activities, and it seems intuitive that proper training would result in higher quality questions. The extent to which this is true would be an interesting avenue for future work.

The tutorials on MCQ criticism were exceptionally easy to run and our students provided positive feedback on these experiences. The tutorials on Bloom’s Taxonomy were also valuable in getting students to recognise that MCQs can test higher-level thinking rather than just recall, and the classes were highly effective at giving them the frames of reference and vocabulary for criticising peers’ questions.

The workload associated with the training was not onerous. The first input from academics is eminently sustainable: the identification and allocation of learning outcomes at suitable granularity, to ensure broad syllabus coverage and to make question selection (Cycle 1) or authoring (Cycle 2) straightforward for students. Using the in-built PeerWise reports and custom dashboards, assessment of these tasks was rapid and communication of feedback easy, encouraging students to give timely, meaningful and actionable feedback to their peers.

## Conclusion

We have demonstrated a practical approach for training students in MCQ design and deploying student-authored questions onto a high-stakes summative assessment. With selection of suitable candidate items resulting from rigorous analysis of performance metrics, the questions provide a defensible assessment of student performance within the unit of study. We found that the selected student-authored MCQs performed as effectively as, and sometimes better than, academic authored and textbook derived MCQs. Leveraging the effort of the student cohort in this fashion represents an opportunity for academics to sustainably build large banks of high performing, syllabus-aligned MCQs with only a modest impact on workload.

